# Dentification of the Sorghum TMCO Gene Family and Differential Expression Analysis Under Aphid Infestation

**DOI:** 10.1101/2025.03.28.646042

**Authors:** Minghui Guan, Chengzhi Ye, Jijuan Li, Shengyun Shan, Tonghan Wang, Yongfei Wang, Lu Sun, Junli Du, Degong Wu

**Author notes:** Correspondence (J. D.). (D. W.).

## Abstract

The three-domain multicopper oxidase (TMCO) family, the largest subfamily of multicopper oxidases, plays a crucial role in plant growth, development, and responses to biotic stresses. To investigate the expression characteristics of the *SbTMCO* gene family under aphid infestation, a genome-wide identification and analysis of TMCO family members in sorghum were conducted using bioinformatics methods. Transcriptome data and qRT-PCR were utilized to preliminarily examine sorghum’s responses to aphid stress. The results revealed 42 *TMCO* genes in the sorghum genome, distributed across 10 chromosomes and classified into five subgroups. Members within the same subgroup displayed similar physicochemical properties, gene structures, and conserved motifs. Collinearity analysis suggested that tandem duplication and segmental duplication were the primary drivers of TMCO gene expansion in sorghum. Cis-acting element analysis identified multiple jasmonic acid (MeJA) and abscisic acid (ABA) responsive elements among *SbTMCO* genes. Gene expression analysis, based on RNA-seq and qRT-PCR, confirmed the relative expression levels of TMCO genes under aphid stress. This study establishes a foundation for exploring the anti-aphid functions of *SbTMCO* genes and provides recommendations and directions for further research on this gene family.

## 1. Introduction

*Sorghum bicolor*, originating from Africa, is an important dual-purpose crop serving as both a food and forage resource(<u>Tanwar, Panghal, Chaudhary, Kumari, & Chhikara, 2023</u>; <u>Winchell, Stevens, Murphy, Champion, & Fuller, 2017</u>). It is a critical source of nutrients and bioactive compounds in human diets. Known as the "camel of crop", sorghum possesses remarkable traits such as high photosynthetic efficiency, drought and waterlogging tolerance, and salinity resistance(<u>Jiahui et al., 2019</u>). Additionally, it serves as an industrial raw material, used for producing beverages and beer(<u>Hariprasanna & Rakshit, 2016</u>). In the Americas, sorghum is primarily cultivated as livestock feed(<u>Dabija, Ciocan, Chetrariu, & Codin, 2021</u>).

The sugarcane aphid (*Melanaphis sacchari*) is a major pest of sorghum species and a key vector for viruses affecting sorghum(<u>Shrestha & Huang, 2022</u>). Since its outbreak in 2013, the sugarcane aphid has transitioned from being a minor to a major pest, posing significant threats to global sorghum production and causing severe economic losses(<u>Esquivel, Faris, & Brewer, 2021</u>). These aphids extract nutrients by piercing sorghum leaves, leading to dehydration and wilting. Prolonged infestation blackens the leaves, reducing photosynthetic efficiency. Affected plants display stunted development, leaf yellowing, necrosis, fewer panicles, and lower thousand-grain weight(<u>Omkar, 2021</u>). Additionally, aphid honeydew secretion promotes the growth of sooty mold, severely impacting sorghum yield and quality(<u>B. U. Singh, Padmaja, & Seetharama, 2004</u>).

Multi-copper Oxidase (MCO) represent a superfamily of genes, encompassing enzymes such as nitrite reductase, laccase, ascorbate oxidase, bilirubin oxidase, ferroxidase, and ceruloplasmin(<u>Mccaig, Meagher, & Dean, 2005</u>; <u>Nakamura, Kawabata, Yura, & Go, 2003</u>; <u>Ye, Pan, Kang, Lu, & Zhang, 2015</u>). MCOs include enzymes with two, three, or six conserved domains, classifying them into two-domain, three-domain, and six-domain MCOs(<u>Chen Huilong 2021,19(17):5569-5580.(in Chinese)</u>). Among these, the three-domain MCOs form the largest subfamily, featuring three conserved domains: Cu-oxidase, Cu-oxidase_2, and Cu-oxidase_3. This family primarily consists of laccases and ascorbate oxidases.

Laccases, first identified in the resin of the Japanese lacquer tree(<u>Khatami et al., 2022</u>), were predominantly studied in fungi for an extended period. Recent years have seen increased research on plant laccases. Studies revealed that deletion of laccase genes in *Cryphonectria parasitica* led to reduced virulence and pathogenicity(<u>Boumnigel, 2021</u>), while laccase gene deficiency in *Sinorhizobium meliloti* resulted in decreased stress resistance and pathogenicity(<u>Anna et al., 2016</u>). Furthermore, research by Li Huifang et al. demonstrated that laccase genes in aphid-susceptible apple varieties like ‘Red Fuji’ exhibited expression patterns opposite to those in resistant varieties, indicating the pivotal role of laccase gene expression in plant anti-aphid responses(<u>LI Huifang 2023, 40(6): 1202-1214. (in Chinese)</u>). Ascorbate oxidase (AAO) is a copper-containing enzyme localized in the apoplast. It catalyzes the oxidation of ascorbate (AA) to dehydroascorbic acid (DHA) through the intermediate monodehydroascorbate (MDHA)(<u>Braunschmid et al.</u>; <u>Xu et al., 2024</u>). Numerous studies indicate that ascorbate enhances plant resistance to biotic stresses through interactions with redox and hormone systems. For instance, Sing et al. demonstrated that AO application in sugar beet roots significantly reduced cyst nematode infestation, decreasing both female and cyst counts compared to untreated plants(<u>R. R. Singh, Nobleza, Demeestere, & Kyndt, 2020</u>). Elevated ascorbate oxidase activity raises the redox ratio of the ascorbate pool, altering the redox gradient across the plasma membrane. In Arabidopsis thaliana, these changes regulate stomatal opening, stress responses, and gene expression, thereby influencing growth and development(<u>Bérczi, Caubergs, & Asard, 2003</u>).

This study utilized aphid-resistant sorghum variety HX133 and aphid-susceptible sorghum variety HX141 to conduct a bioinformatics analysis of the three-domain multicopper oxidase (TMCO) gene family. The research focused on examining the structure of TMCO family proteins and the phylogenetic relationships among family members. Key analyses included the identification of family members, phylogenetic evolution, gene structure and conserved motifs, and promoter cis-acting regulatory elements. Additionally, the expression of TMCO genes under aphid stress was investigated using qRT-PCR to determine their expression levels at different time points during aphid infestation. The findings aim to elucidate the role of TMCO genes in sorghum’s resistance to aphids, providing new insights into the function of multicopper oxidases in plant defense against insect pests.

## 2. Materials and Methods

### 2.1 Identification and Retrieval of *SbTMCO* Gene Family

The genome sequence and annotation files of sorghum were downloaded fr om the NCBI database (Sorghum_bicolor_NCBIv3). Using the Gtf/Gff3 sequence extractor tool in the TBtools software, the CDS sequences of all sorghum genes were extracted from the genome. The hmm model of the *SbTMCO* family (Pfam ID: PF00394) was obtained from the Pfam database (http://pfam.xfam.org/)(<u>Finn et al., 2014</u>). GeneDoc software was employed to align the selected amino acid sequences of *SbTMCO*s(<u>Nicholas</u>). Finally, the online SMART tool (https://smart.embl.de/) was used to verify the presence of the conserved Cu-oxidase, Cu-oxidase_2, and Cu-oxidase_3 domains, confirming the identified sequences a s TMCO family members(<u>Ivica & Peer, 2018</u>).

### 2.2 Physicochemical Properties and Subcellular Localization of *SbTMCO* Family

The physicochemical properties of the *SbTMCO* family members, such as amino acid count, isoelectric point (pI), and molecular weight (MW), were analyzed using the ExPASy ProtParam tool (https://web.expasy.org/protparam/). Subc ellular localization predictions were performed with the WoLF PSORT platform (https://wolf-psort.hgc.jp/). Additionally, protein secondary structures were predicted using the SOPMA tool (https://npsa-prabi.ibcp.fr/cgi-bin/npsa_automat.pl?page=npsa_sopma.html)(Duvaud, Gabella, Lisacek, Stockinger, & Durinx, 2021; Paul et al., 2007).

### 2.3 Chromosomal Localization, Collinearity, and Ka/Ks Analysis of *SbTMCO* Family

The sorghum genome and annotation files downloaded from NCBI were processed using the Fasta Stats tool in the TBtools software suite to generate chromosome length files. Gene duplication relationships were analyzed using the One Step MCScanX plugin in TBtools. The related gene pairs were then input into the Simple Ka/Ks Calculator plugin in TBtools to compute the Ka and Ks values. The divergence time was calculated using the formula T=Ks/(2λ)( λ=6.5×10^-9^)(<u>Chen, Wu, & Li, 2023</u>).

### 2.4 Gene Structure, Conserved Motif Analysis, and *Cis*-Acting Element Prediction of *SbTMCO* Family

The exon, intron, and UTR information of *SbTMCO* genes was extracted from the genome annotation file using TBtools software v2.042. Conserved motifs were detected using the MEME Suite (https://meme-suite.org/meme/), identifying the 10 most conserved motifs in *SbTMCO* proteins(<u>Bailey et al., 2009</u>). Chromosomal distribution data for the TMCO family were obtained through the genome annotation file. Finally, TBtools software v2.042 was used to visualize gene structures, conserved motifs, and cis-acting element information(<u>Lescot & M., 2002</u>).

### 2.5 Collinearity Analysis and Phylogenetic Tree Construction of *SbTMCO* Family

Collinearity relationships between sorghum, soybean, and Arabidopsis were analyzed using the **One Step MCScanX** tool in TBtools(<u>Yupeng et al., 2012</u>) (default parameters). A phylogenetic tree was constructed using the Neighbor-Joining (NJ) method in MEGA11, encompassing 42 three-domain multicopper oxidase protein sequences. Optimization of the phylogenetic tree was conducted using the "One Step Build a ML Tree" tool in TBtools. Collinearity maps were generated with the **Multiple Synteny Plot** tool in TBtools and visually enhanced using Adobe Illustrator (2023).

### 2.6 Transcriptomic Analysis of *SbTMCO* Genes

Transcriptome data under biotic stress conditions were downloaded from the NCBI database to assess *SbTMCO* expression under aphid infestation (PRJNA716317). Data quality was ensured using the FastQC tool (https://github.com/s-andrews/FastQC). Filtered data were processed with the **Kallisto Super Wrapper V3** tool in TBtools. Following Zou’s methodology(<u>Zou et al., 2024</u>), the expression of TMCO family members was analyzed under aphid stress conditions (6h, 24h, 48h, 7d). Differential expression heatmaps were visualized using TBtools software v2.042.

### 2.7 Experimental Materials and Aphid Rearing

Aphids were collected from sorghum plants in the western planting area of Anhui Science and Technology University and reared in an artificial climate chamber (26 ± 1°C, 50% relative humidity, 16 h light/8 h dark photoperiod). The population was maintained by transferring aphids to fresh plants biweekly.

This study utilized resistant sorghum variety HX133 and susceptible variety HX141. Mature, non-dormant seeds were sown in pots containing a mixture of nutrient soil and vermiculite. Plants were cultivated in an artificial climate chamber under conditions of 26 ± 1°C, 50% relative humidity, and a 16 h light/8 h dark cycle. At the three-leaf stage, experiments were initiated. Three seedlings were harvested per time point as biological replicates. Samples were flash-frozen in liquid nitrogen upon collection and stored at −80 °C to prevent RNA degradation.

### 2.8 RNA Extraction and qRT-PCR Validation

Total RNA was extracted from sorghum samples using the Trizol method. The concentration and quality of the extracted RNA were evaluated with an ultra-low-volume spectrophotometer by measuring the A260/A280 ratio, ensuring values fell within the range of 1.8–2.1. High-quality RNA samples were then reverse-transcribed into cDNA using HiScript QRT reagents. The resulting cDNA was diluted fivefold for qRT-PCR validation, with each sample analyzed in triplicate.

Relative expression levels of the target genes were calculated using the 2^−ΔΔCt^ method. Statistical analyses were performed using SPSS 24.0 software, and bar charts were generated with GraphPad Prism 8(<u>Livak & Schmittgen, 2001</u>; <u>Motulsky, 2007</u>). This methodology ensures the accuracy of RNA quality checks and provides reliable data for analyzing gene expression under experimental conditions.

## 3. Results and Analysis

### 3.1 Chromosomal Localization of *SbTMCO* Family Genes

Based on the chromosomal length information and annotation file, chromosomal mapping revealed that the 42 TMCO genes were distributed randomly across 12 chromosomes (Figure 1). A detailed distribution analysis showed that chromosomes 1, 2, and 3 contained the majority of the TMCO family members (57%), while chromosomes 4, 5, 6, and 10 had four TMCO genes each. Chromosomes 6, 7, and 8 had only one TMCO gene, whereas chromosome 9 contained three genes. Interestingly, genes located on chromosomes 3, 4, and 9 were clustered or duplicated in specific regions, indicating the presence of gene clusters or segmental duplications. These patterns suggest potential roles in gene expression regulation, functional specificity, and genetic evolution.

**Figure. 1.**
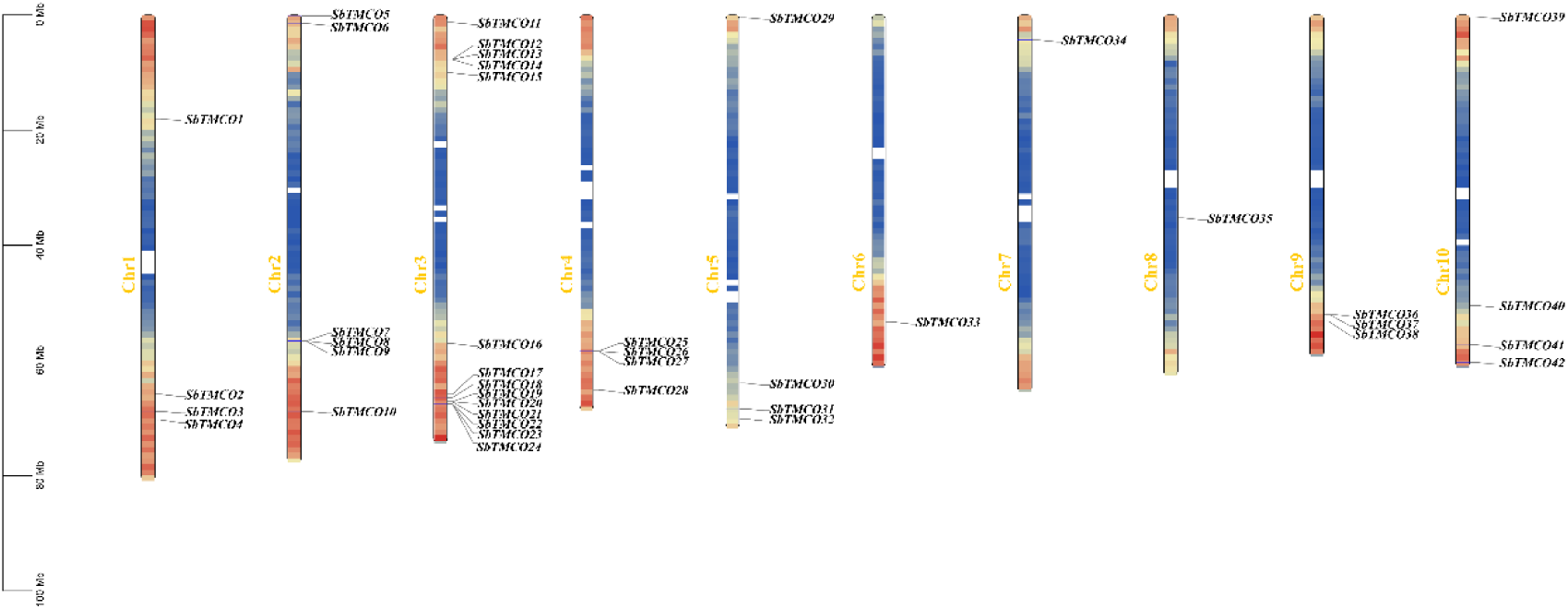
Chromosomal localization and gene duplication analysis of the *SbTMCO* gene

### 3.2 Physiochemical Properties and Subcellular Localization Prediction of *SbTMCO* Family

An analysis of physiochemical properties (Table 1) showed that the amino acid lengths of TMCO proteins ranged from 471 to 949 residues, with molecular weights directly proportional to their lengths. The grand average of hydropathy (GRAVY) analysis revealed that most TMCO proteins were hydrophilic (negative GRAVY values), while proteins such as *SbTMCO3*, *SbTMCO19*, *SbTMCO24*, *SbTMCO20*, *SbTMCO4*, and *SbTMCO36* were hydrophobic (positive GRAVY values).

**Table 1.** Analysis of physicochemical properties and subcellular localization of the TMCO gene family in sorghum.

| Gene name | Number of<br>Amino Acid | Molecular<br>Weight | Theoretical<br>pI | Instability<br>Index | Aliphatic<br>Index | Average<br>coefficient of<br>hydrophilicity | Subcellular<br>localization |
| --- | --- | --- | --- | --- | --- | --- | --- |
| *SbTMCO1* | 599 | 66343.66 | 8.2 | 37.52 | 81.82 | -0.292 | vacu |
| *SbTMCO2* | 949 | 105606.97 | 8.45 | 35.32 | 86.77 | -0.169 | plas |
| *SbTMCO3* | 600 | 64954.6 | 7.02 | 41.71 | 85.2 | 0.077 | mito |
| *SbTMCO4* | 576 | 62694.72 | 8.49 | 31.59 | 83.28 | 0.002 | chlo |
| *SbTMCO5* | 542 | 58723.74 | 7.34 | 33.73 | 80.31 | -0.044 | cyto |
| *SbTMCO6* | 553 | 59428.84 | 5.6 | 37.64 | 86.69 | -0.058 | cyto |
| *SbTMCO7* | 570 | 62655.01 | 7.09 | 36.61 | 76.65 | -0.219 | chlo |
| *SbTMCO8* | 571 | 62665.21 | 8.03 | 32.25 | 79.77 | -0.181 | vacu |
| *SbTMCO9* | 579 | 63460.87 | 6.24 | 33.29 | 79.52 | -0.187 | chlo |
| *SbTMCO10* | 548 | 60502.51 | 9.34 | 37.86 | 82.9 | -0.178 | cyto |
| *SbTMCO11* | 587 | 64375.01 | 6.8 | 41.82 | 82.37 | -0.191 | extr |
| *SbTMCO12* | 588 | 64318.46 | 5.97 | 41.18 | 86.31 | -0.051 | vacu |
| *SbTMCO13* | 591 | 65420.45 | 6.92 | 46.04 | 76.23 | -0.276 | extr |
| *SbTMCO14* | 610 | 67735.18 | 6.29 | 41.36 | 81.56 | -0.204 | plas |
| *SbTMCO15* | 598 | 66222.55 | 6.31 | 36.94 | 76.94 | -0.182 | vacu |
| *SbTMCO16* | 471 | 51433.75 | 8.86 | 39.69 | 83.65 | -0.182 | cyto |
| *SbTMCO17* | 558 | 62004.87 | 8.78 | 36.67 | 78.46 | -0.24 | vacu |
| *SbTMCO18* | 568 | 62830.71 | 6.01 | 33.23 | 83.75 | -0.063 | extr |
| *SbTMCO19* | 579 | 63139.54 | 7.67 | 28.22 | 90.43 | 0.076 | extr |
| *SbTMCO20* | 579 | 63104.31 | 8.02 | 32.2 | 86.68 | 0.046 | extr |
| *SbTMCO21* | 579 | 63559.19 | 8.5 | 34.42 | 78.81 | -0.155 | chlo |
| *SbTMCO22* | 496 | 54243.19 | 6.57 | 42.27 | 72.04 | -0.194 | mito |
| *SbTMCO23* | 649 | 68089.36 | 5.88 | 41.58 | 83.57 | 0.12 | plas |
| *SbTMCO24* | 557 | 60246.15 | 6.49 | 39.56 | 81.8 | 0.068 | mito |
| *SbTMCO25* | 591 | 65427.05 | 6.38 | 37.91 | 77.39 | -0.232 | chlo |
| *SbTMCO26* | 590 | 65422.48 | 6.06 | 33.74 | 75.98 | -0.215 | cyto |
| *SbTMCO27* | 573 | 63909.64 | 6.23 | 35.07 | 72.84 | -0.261 | chlo |
| *SbTMCO28* | 591 | 65441.75 | 7.08 | 34.76 | 77.01 | -0.182 | chlo |
| *SbTMCO29* | 587 | 63243.46 | 5.83 | 32.29 | 78.94 | -0.013 | chlo |
| *SbTMCO30* | 574 | 63292.69 | 5.75 | 37.48 | 79.44 | -0.209 | vacu |
| *SbTMCO31* | 601 | 66657.79 | 6.02 | 38.41 | 80.2 | -0.195 | extr |
| *SbTMCO32* | 600 | 65464.03 | 5.75 | 34.47 | 79.38 | -0.146 | extr |
| *SbTMCO33* | 578 | 63063.91 | 6.57 | 37.69 | 80.67 | -0.214 | chlo |
| *SbTMCO34* | 505 | 56768.59 | 6.46 | 34.1 | 73.37 | -0.44 | cyto |
| *SbTMCO35* | 585 | 64259.54 | 5.08 | 33.25 | 80.32 | -0.089 | vacu |
| *SbTMCO36* | 585 | 63385.43 | 9.32 | 37.33 | 84.19 | 0.001 | chlo |
| *SbTMCO37* | 585 | 63239.71 | 5.95 | 35.37 | 82.96 | -0.01 | vacu |
| *SbTMCO38* | 559 | 62219.84 | 7.29 | 39.62 | 81.77 | -0.207 | vacu |
| *SbTMCO39* | 590 | 65565.24 | 5.84 | 30.87 | 80.76 | -0.238 | plas |
| *SbTMCO40* | 638 | 70042.9 | 5.93 | 41.1 | 76.77 | -0.278 | chlo |
| *SbTMCO41* | 546 | 60697.76 | 8.6 | 43.35 | 83.57 | -0.187 | chlo |
| *SbTMCO42* | 605 | 66303.62 | 9.08 | 35.03 | 79.17 | -0.145 | chlo |

Note: chlo is chloroplast; cyto is cytoplasm; vacu is vesicle; extr is extracellular matrix; plas is plasma membrane; mito is mitochondria

Among the proteins, 11 were acidic (pI < 6.5), 22 were basic (pI > 7.5), and 9 were neutral (pI = 6.5–7.5). The highest isoelectric point (pI) was 9.43, while the lowest was 5.08. The instability index ranged from 28.22 to 46.94, and the aliphatic index varied from 72.04 to 90.43, indicating diverse structural properties within the family.

Subcellular localization prediction revealed that most TMCO proteins were targeted to chloroplasts, vacuoles, and the extracellular matrix, with a few localized in the cytoplasm. This distribution highlights the functional diversity of TMCO proteins in sorghum.

Secondary structure analysis revealed that all *SbTMCO* proteins are composed of α-helices, extended strands, and random coils (Table 2). The proportion of α-helices ranged from 5.08% to 21.81%, with *SbTMCO* having the highest proportion and *SbTMCO28* the lowest. Extended strands accounted for 20.82% to 29.54%, while random coils constituted 54.27% to 72.79%, making them the dominant structural element.These variations in secondary structure suggest significant spatial differences among *SbTMCO* proteins, potentially reflecting diverse biological functions.

**Table 2.** Secondary structure prediction of the *SbTMCO* gene family.

| Gene name | Alpha helix | Extended strand | Random coil |
| --- | --- | --- | --- |
| *SbTMCO1* | 63(10.52%) | 152(25.38%) | 384(64.11s%) |
| *SbTMCO2* | 207(21.81%) | 227(23.92%) | 515(54.27%) |
| *SbTMCO3* | 70(11.69%) | 150(25.04%) | 379(63.27%) |
| *SbTMCO4* | 51(8.85%) | 150(26.04%) | 375(65.1%) |
| *SbTMCO5* | 37(6.81%) | 147(27.07%) | 359(66.11%) |
| *SbTMCO6* | 50(9.04%) | 157(28.39%) | 342(62.57%) |
| *SbTMCO7* | 49(8.6%) | 149(26.14%) | 372(65.26%) |
| *SbTMCO8* | 33(5.78%) | 147(25.74%) | 391(68.48%) |
| *SbTMCO9* | 63(10.88%) | 146(25.22%) | 370(63.9%) |
| *SbTMCO10* | 67(12.23%) | 149(27.19%) | 332(60.58%) |
| *SbTMCO11* | 52(8.86%) | 145(24.7%) | 390(66.44%) |
| *SbTMCO12* | 51(8.67%) | 129(21.94%) | 408(69.39%) |
| *SbTMCO13* | 57(9.64%) | 128(21.66%) | 406(68.7%) |
| *SbTMCO14* | 39(6.39%) | 127(20.82%) | 444(72.79%) |
| *SbTMCO15* | 63(10.54%) | 145(24.25%) | 390(65.22%) |
| *SbTMCO16* | 29(6.12%) | 140(29.54%) | 305(64.35%) |
| *SbTMCO17* | 60(10.75%) | 148(26.52%) | 350(62.72%) |
| *SbTMCO18* | 52(9.5%) | 149(26.23%) | 367(64.61%) |
| *SbTMCO19* | 53(9.15%) | 148(25.56%) | 378(65.28%) |
| *SbTMCO20* | 67(11.57%) | 153(26.42%) | 359(62%) |
| *SbTMCO21* | 62(10.71%) | 154(26.6%) | 363(62.92%) |
| *SbTMCO22* | 36(7.26%) | 131(26.41%) | 329(66.33%) |
| *SbTMCO23* | 64(9.86%) | 148(22.8%) | 437(67.33%) |
| *SbTMCO24* | 53(9.52%) | 144(25.82%) | 360(64.43%) |
| *SbTMCO25* | 51(8.63%) | 155(26.23%) | 385(65.14%) |
| *SbTMCO26* | 65(11.02%) | 149(25.25%) | 376(63.73%) |
| *SbTMCO27* | 54(9.42%) | 147(25.65%) | 372(64.92%) |
| *SbTMCO28* | 30(5.08%) | 152(25.76%) | 408(69.15%) |
| *SbTMCO29* | 59(10.05%) | 145(24.7%) | 383(65.25%) |
| *SbTMCO30* | 42(7.32%) | 151(26.31%) | 381(66.38%) |
| *SbTMCO31* | 56(9.32%) | 146(24.29%) | 399(66.39%) |
| *SbTMCO32* | 48(8%) | 154(25.67%) | 398(66.33%) |
| *SbTMCO33* | 58(10.03%) | 149(25.78%) | 371(64.19%) |
| *SbTMCO34* | 59(11.68%) | 124(24.55%) | 322(63.76%) |
| *SbTMCO35* | 59(10.09%) | 156(26.67%) | 370(63.25%) |
| *SbTMCO36* | 46(7.86%) | 157(26.84%) | 382(65.3%) |
| *SbTMCO37* | 52(8.89%) | 151(25.81%) | 382(65.3%) |
| *SbTMCO38* | 68(12.16%) | 142(25.4%) | 349(62.43%) |
| *SbTMCO39* | 76(12.88%) | 156(26.44%) | 358(60.68%) |
| *SbTMCO40* | 57(8.93%) | 144(22.57%) | 437(68.5%) |
| *SbTMCO41* | 56(10.26%) | 152(27.48%) | 338(61.9%) |
| *SbTMCO42* | 58(9.59%) | 152(25.12%) | 395(65.29%) |

### 3.3 Phylogenetic Analysis of *SbTMCO* Family Proteins

To better understand the evolutionary relationships among *SbTMCO* proteins, a phylogenetic tree was constructed using TMCO protein sequences from sorghum, *Arabidopsis thaliana*, and soybean (Figure 2). Based on clustering, the *SbTMCO* proteins were grouped and named according to the branches. The phylogenetic analysis revealed five clades: Clade I contained 57 members, making it the largest group; Clade III included 56 members; Clade II had 5 members; Clade IV contained 35 members; Clade V included 13 members.

**Figure. 2.**
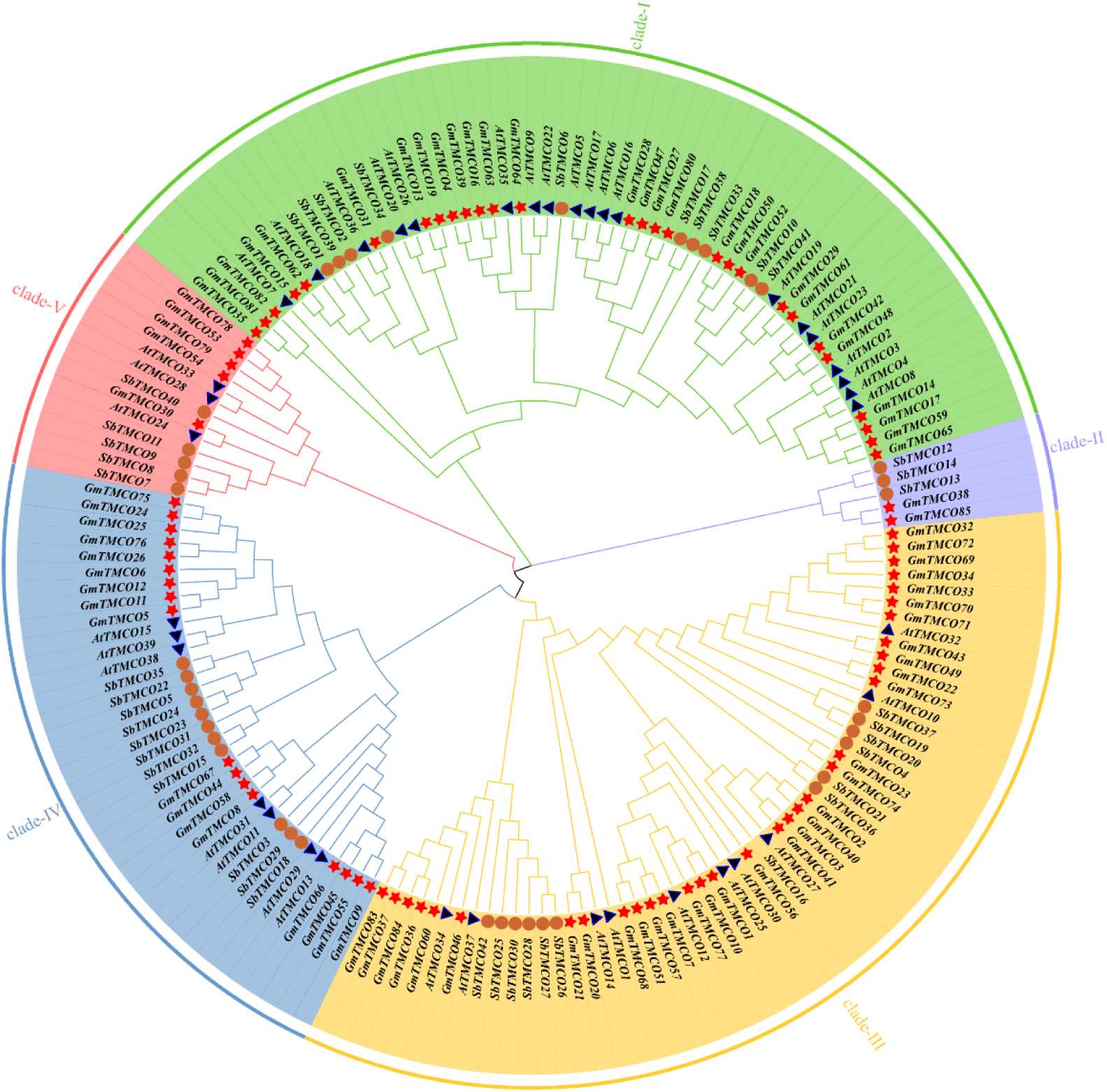
phylogenetic association of plant CCO family proteins Arabidopsis thaliana in blue; soybean in red; sorghum in brown

The distribution of TMCO genes from sorghum, soybean, and *Arabidopsis* was uneven across all clades, indicating evolutionary divergence and functional diversification of the TMCO gene family across species.

### 3.4 Gene Structure, Functional Domain, and Conserved Motif Analysis of the *SbTMCO* Family

The conserved motif analysis revealed that the *SbTMCO* gene family contains ten conserved motifs (Figure 3). Among them, *SbTMCO12*, *SbTMCO13*, and *SbTMCO14* possess only four conserved motifs—motifs 6, 4, 7, and 8. These motifs exhibit high conservation, suggesting potential motif loss during gene duplication. Genes within the same subgroup shared similar gene structures and intron-exon distributions, indicating that members within subfamilies might perform similar functions.

**Figure. 3.**
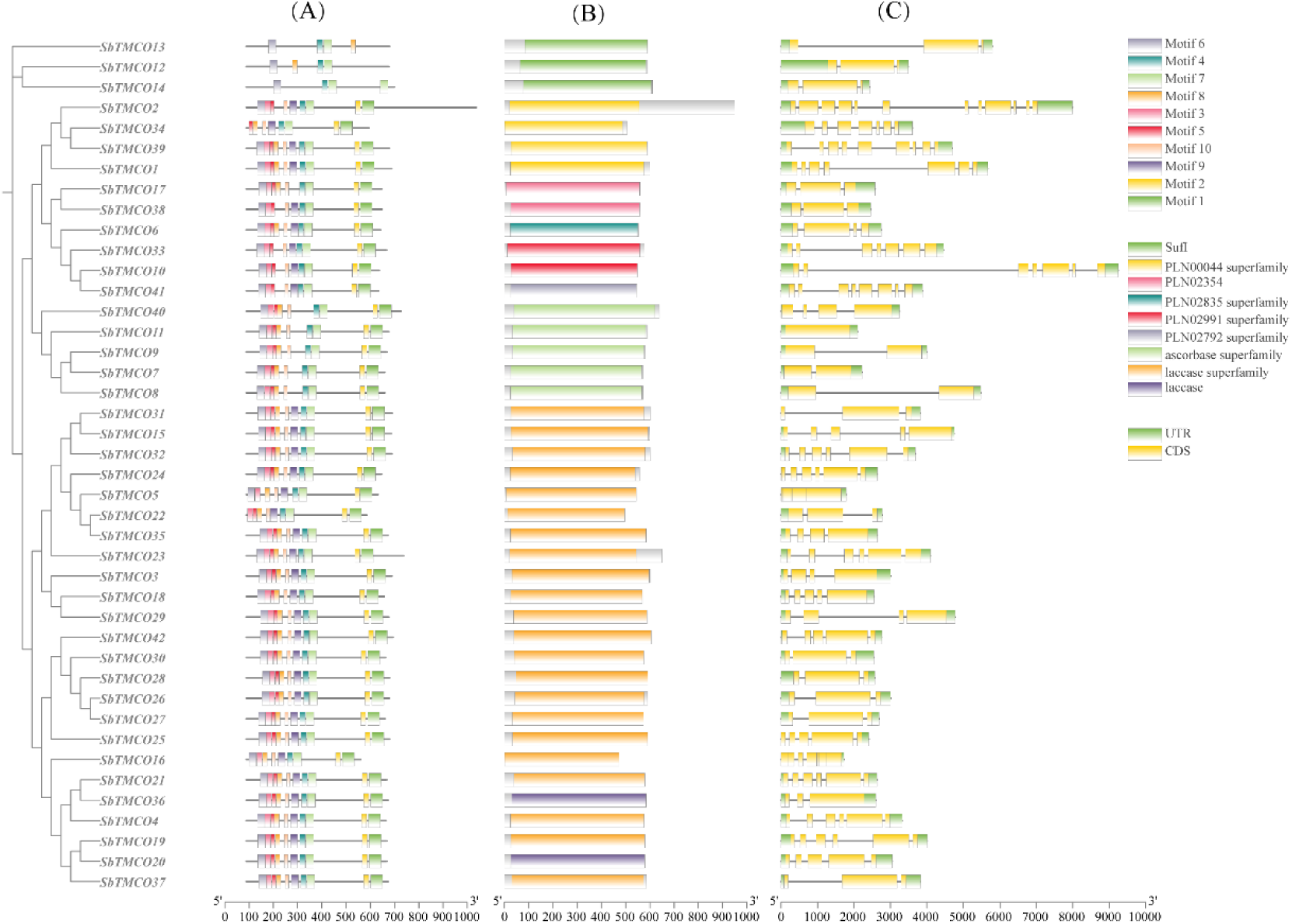
phylogenetic tree, conserved motifs, gene structure and conserved structural domains of the *SbTMCO* gene family

Functional domain analysis identified nine domain types in the *SbTMCO* family: PLN00044, PLN02835, PLN02991, PLN02792, ascorbase, laccase superfamily, SufI, PLN02354, and laccase. Protein motif analysis using MEME uncovered similar patterns, with the order and distribution of motifs generally consistent among subfamily members. Overall, structural domain similarities imply that these genes might share similar functions. However, variations in gene structure and motifs among subfamily members might account for differences in their encoded enzymatic activities.

As shown in Figure 3, the exon numbers in the *SbTMCO* gene family range from 3 to 7. Notably, *SbTMCO12*, *SbTMCO13*, and *SbTMCO14* have simpler gene structures, each containing only two UTRs and three CDSs. *SbTMCO2*, on the other hand, has the highest exon count, with 12 exons. Variations in exon distribution and count may lead to functional differences in genetic engineering and bioinformatics studies. Understanding UTR and CDS composition is crucial for insights into gene expression regulation, gene editing, and protein function research.

### 3.5 Cis-Regulatory Element Analysis of the Promoter Regions in the *SbTMCO* Gene Family

*Cis*-regulatory elements, non-coding sequences in gene promoter regions, play essential roles in regulating the transcription of associated genes. Using the PlantCARE database, regulatory elements in the promoter regions of *SbTMCO* gene family members were identified. A total of 13 cis-regulatory element types were detected (Figure 4). These include regulatory elements related to plant hormone responses— such as methyl jasmonate (MeJ), abscisic acid (ABA), gibberellic acid (GA), salicylic acid (SA), auxin, and flavonoid response elements—and elements associated with abiotic stress, including those involved in defense and stress responses as well as low-temperature responsiveness.

**Figure. 4.**
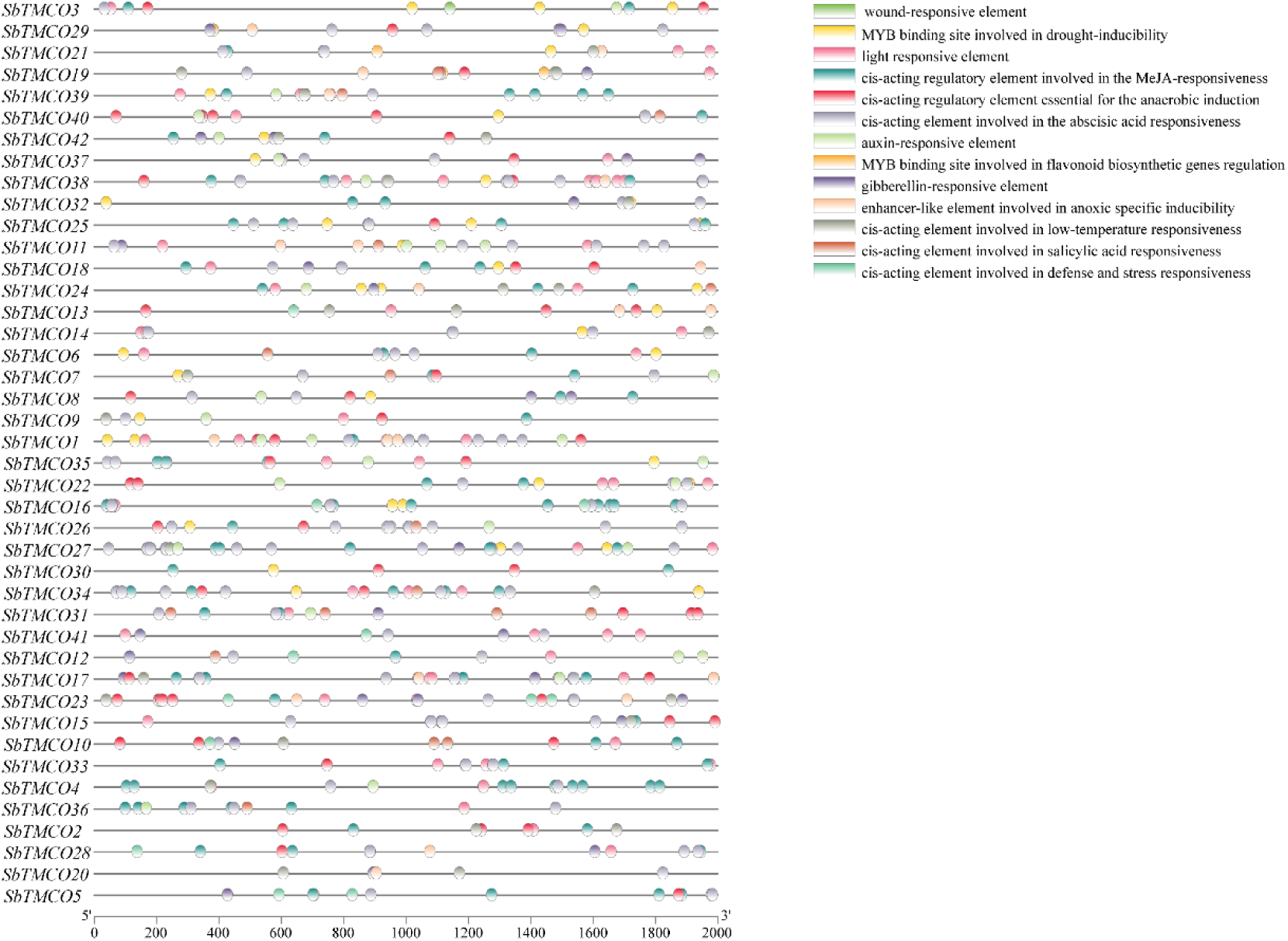
Analysis of cis-elements in the promoter region of the *SbTMCO* gene

Among these, the two most common elements were MeJA-responsive and ABA-responsive elements. These findings suggest that TMCO genes may play crucial roles in responding to aphid-induced stress, providing a theoretical basis for future functional validation of these genes.

### 3.6 Collinearity Analysis and Homologous Evolution Relationships of the *SbTMCO* Gene Family Across Species, and Ka/Ks Analysis

In the sorghum genome, collinearity was identified only between genes located on chromosomes 1, 3, 5, and 9. The types of collinear relationships included whole-genome duplications (WGD) or segmental duplications on chromosomes. The *SbTMCO* family exhibited eight tandem gene duplication events and four segmental duplication events. Among the 12 collinear gene pairs analyzed, the Ka/Ks ratios were all less than 1 (Table 3), indicating a negative selection effect, which implies that these genes are under purifying selection during evolution.

**Table 3.**
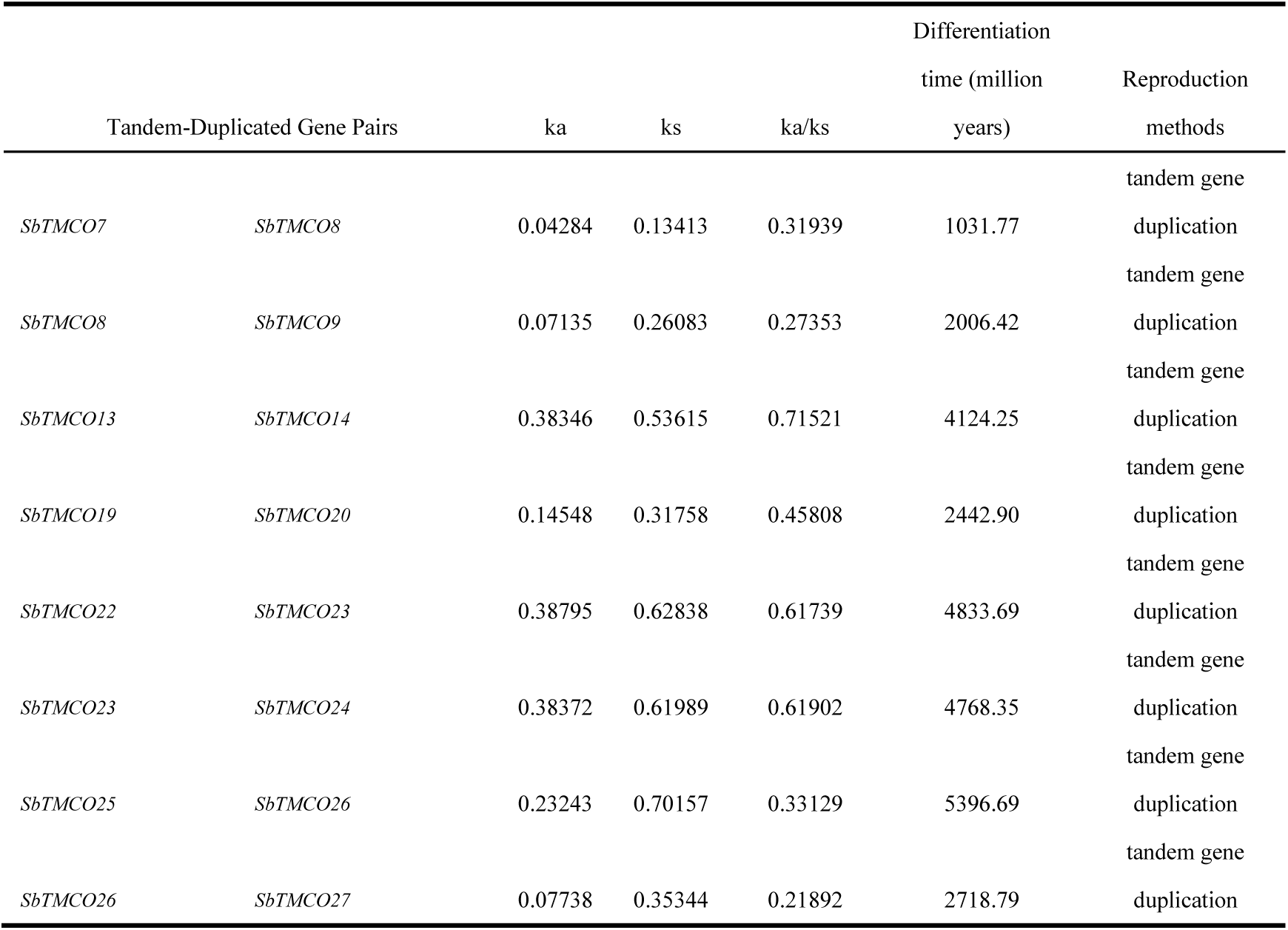
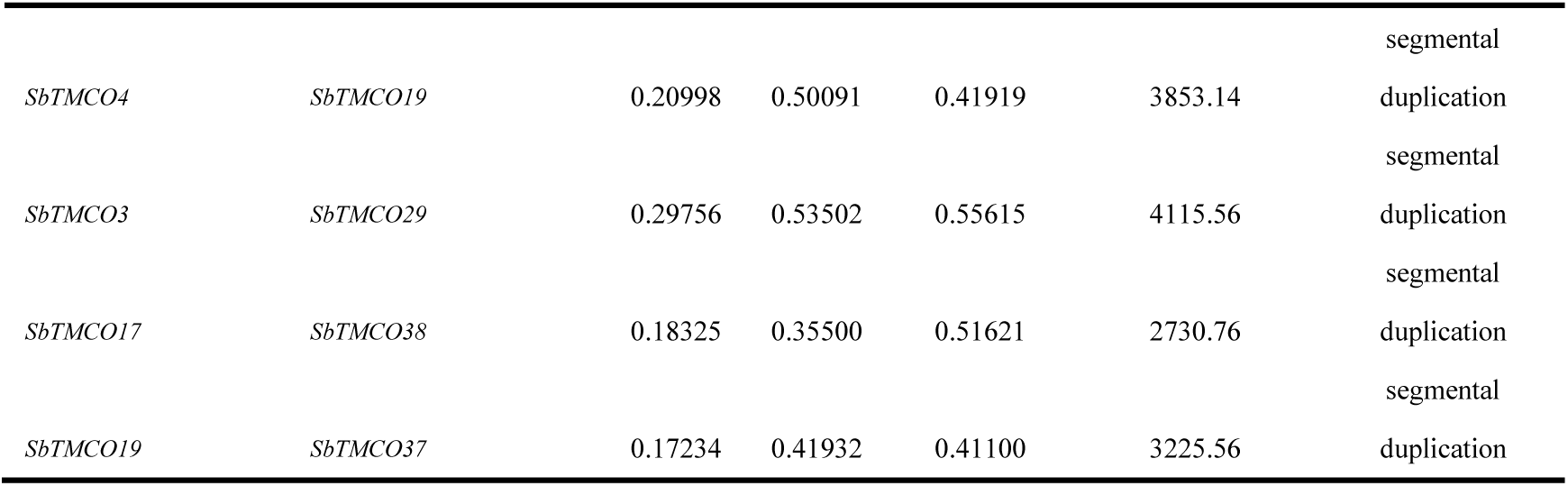
Analysis of *SbTMCO* gene duplication and Ka/Ks values.

**Figure. 5.**
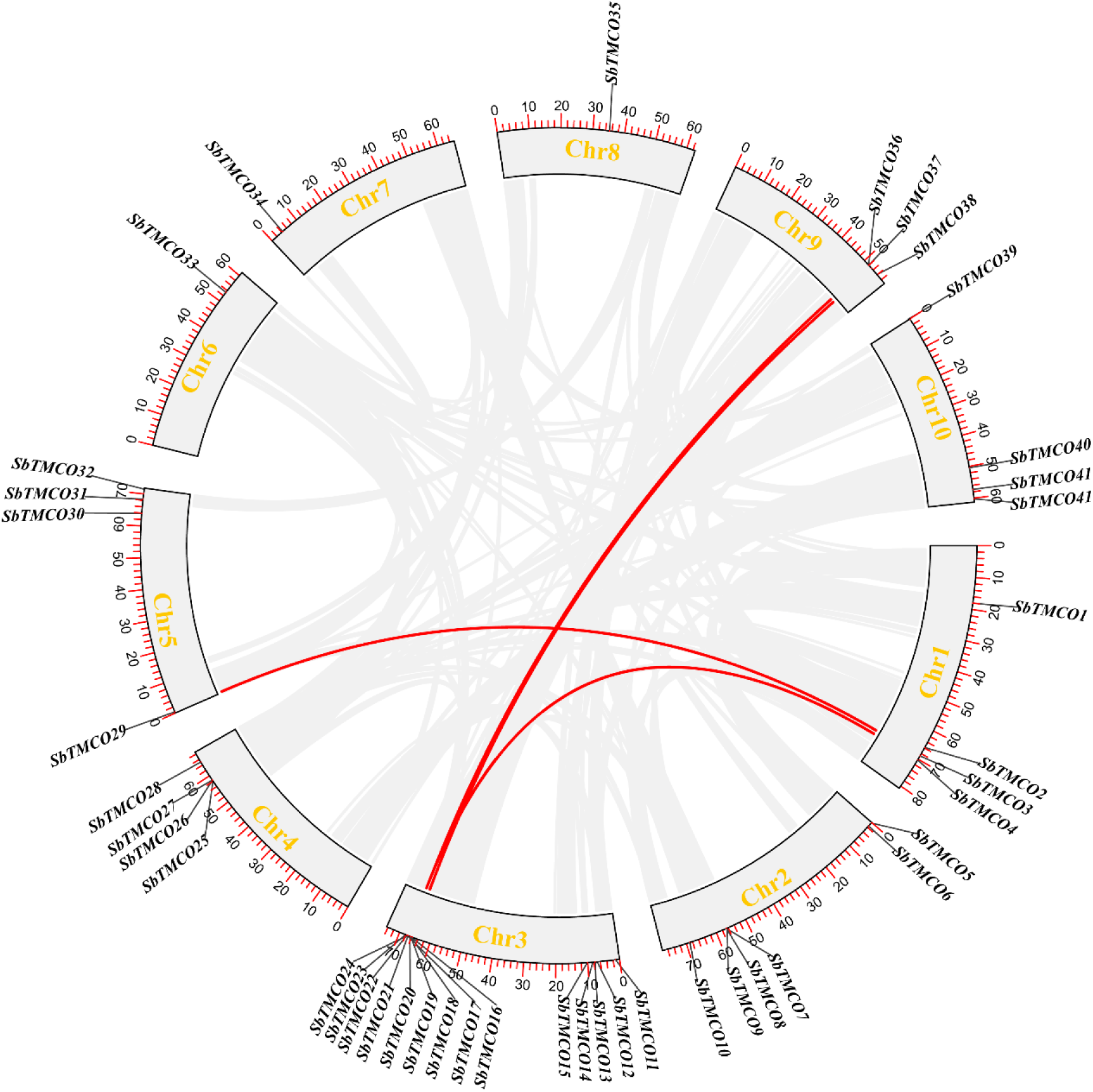
analysis of covariance within the *SbTMCO* family

Through comparative collinearity analysis of *SbTMCO* family members with those of soybean and *Arabidopsis thaliana* (Figure 6), it was observed that 43 collinear gene pairs were shared between sorghum and soybean, whereas 12 collinear pairs were found between sorghum and *Arabidopsis*. Interestingly, *SbTMCO35* did not show collinearity with genes in either of the other two species. The higher number of homologous gene pairs between sorghum and soybean compared to sorghum and *Arabidopsis* suggests that sorghum shares a closer evolutionary relationship with soybean. These interspecies comparisons provide valuable insights into the genetic relationships and functional roles of TMCO genes.

**Figure. 6.**
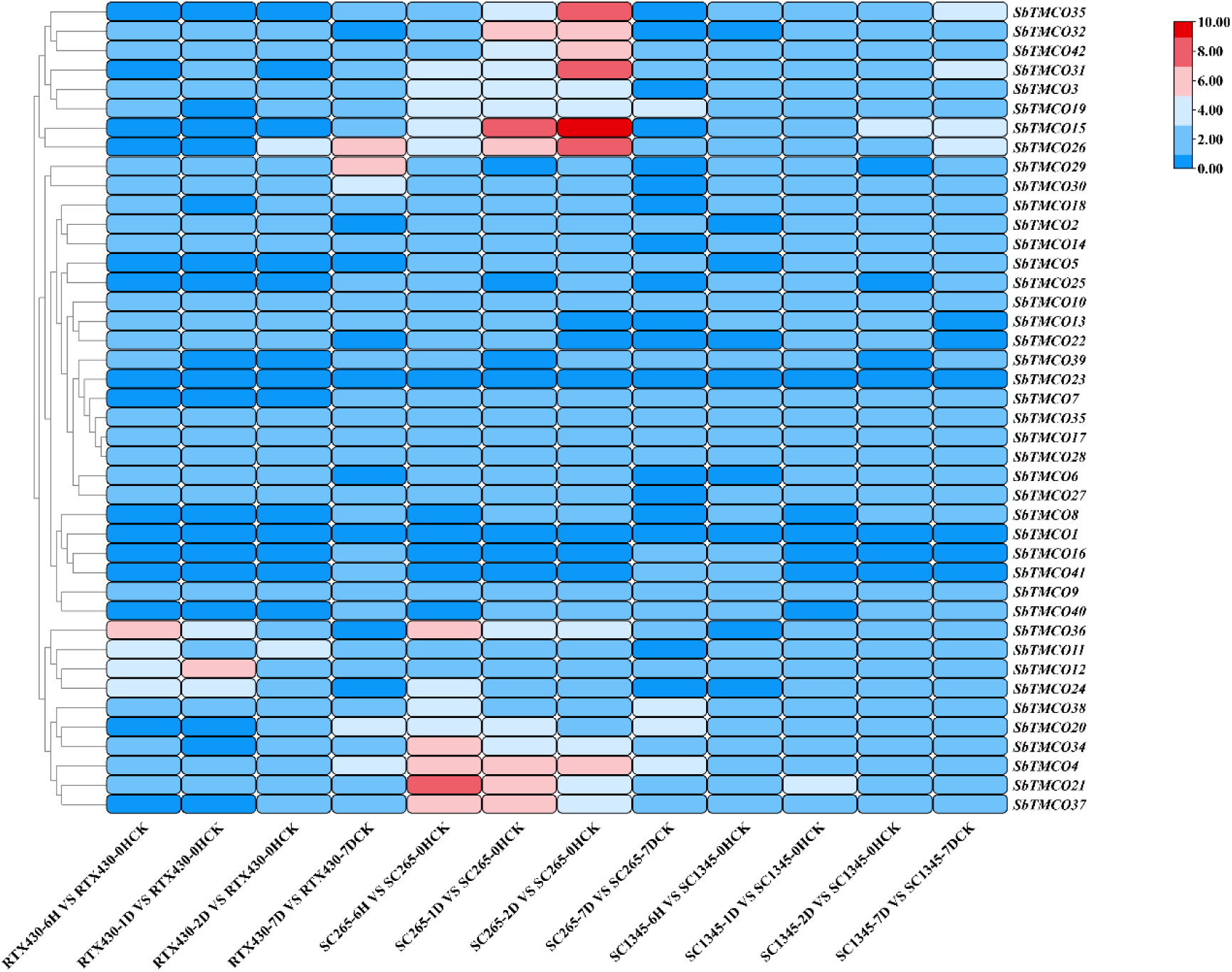
Differential Expression of the Transcriptome of the TMCO Gene Family in Sorghum under Aphid Infestation RTx430 was used as the reference sorghum, SC265 as the resistant sorghum and SC1345 as the susceptible sorghum. Samples were taken at early time points (6h, 24h, 48h) and late time points (7d) after aphid infestation and on 0h and 7d aphid-uninfested plants as controls. Genes with high expression levels are marked with red color.

**Figure. 7.**
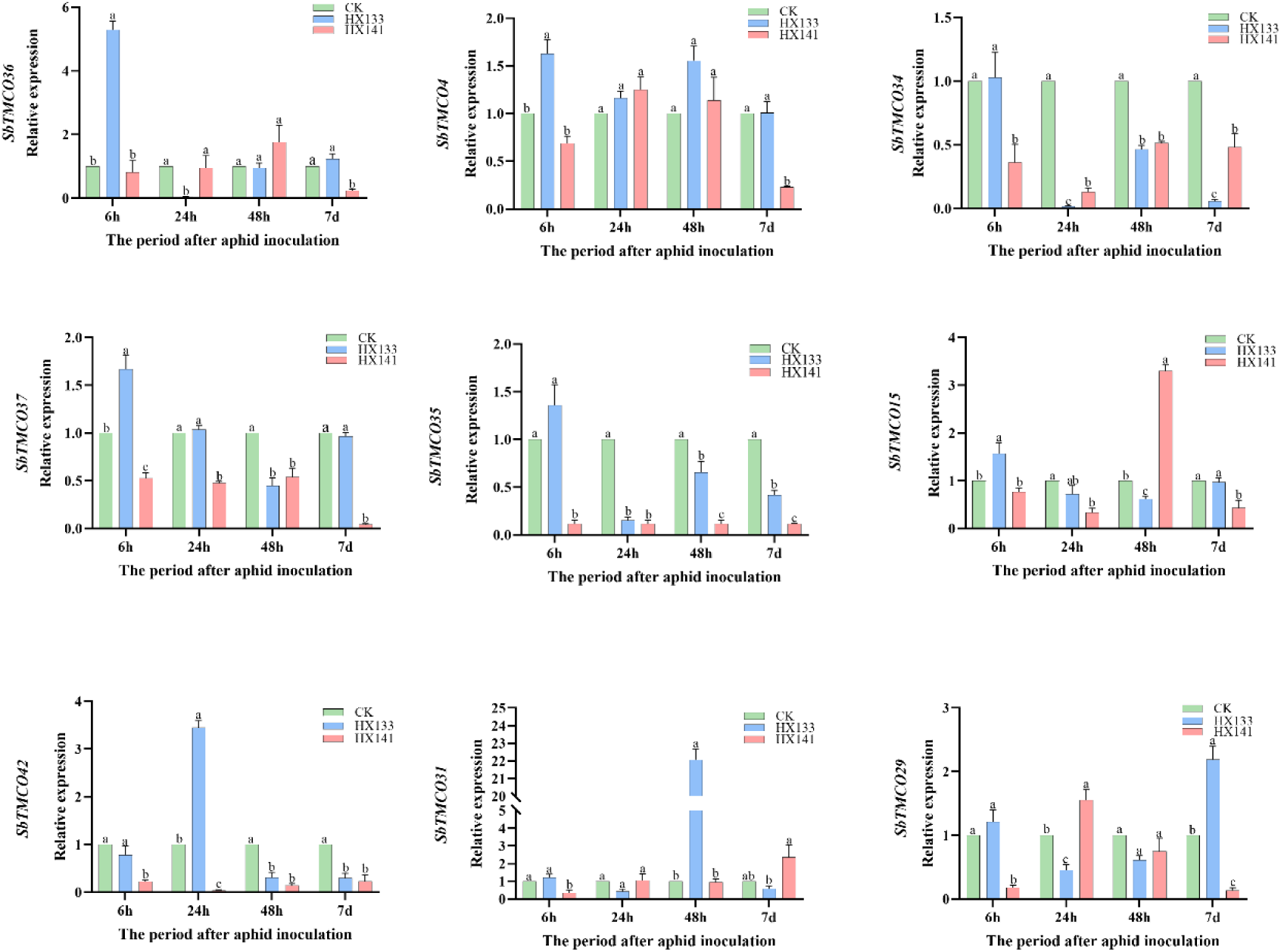
Expression pattern of *SbTMCO* gene family after aphid infestation: 0 h after infestation as early control and 7 days after infestation as late control. Significant differences are indicated by lowercase letters (P < 0.05)

### 3.7 Transcriptome Analysis of the Sorghum MCO Gene Family

Transcriptome analysis revealed distinct expression patterns of *SbTMCO* genes in response to aphid infestation among different sorghum varieties (Figure 8). In aphid-resistant sorghum lines, specific TMCO genes showed significant expression changes, suggesting their roles in conferring resistance. Notably, the highest expression levels of *SbTMCO34*, *SbTMCO4* and *SbTMCO21* and *SbTMCO37* were observed after 6 h of aphid stress. At 1 day (1d), maximum expression of *SbTMCO32*, *SbTMCO15*, *SbTCO26*, *SbSTMCO4*, *SbTMO21* and *SbTMCO7* was observed. At 2 days (2d): significant expression of *SbTMCO35*, *SbTMCO22*, *SbTCO42*, *SbTMCO31*, *SbTMCO15*, *SbSTMCO26* and *SbTMCO4*.

In contrast, expression patterns in aphid-susceptible sorghum varieties were less pronounced. These results highlight the potential roles of specific TMCO genes in mediating sorghum’s resistance mechanisms against aphid-induced stress. Further research is needed to validate these findings and elucidate the underlying molecular mechanisms.

### 3.8 qRT-PCR Expression Analysis of *SbTMCO* Genes Under Aphid Stress

To investigate the expression of *SbTMCO* gene family members during different time points of aphid stress, qRT-PCR was conducted on a resistant sorghum variety, HX133, and a susceptible variety, HX141. The results revealed distinct expression patterns among the TMCO genes (Figure 8): *SbTMCO36*, *SbTMCO4*, *SbTMCO34*, *SbTMCO37*, *SbTMCO35*, *SbTMCO15*, and *SbTMCO29*showed the highest expression at 6h, with *SbTMCO4*, *SbTMCO34*, and *SbTMCO37*showed similar expression profiles to the RNA-seq data. *SbTMCO36*, *SbTMCO35*, and *SbTMCO15* showed different expression profiles; *SbTMCO42* and *SbTMCO31* showed the highest expression at 24h and 48h, respectively; *SbTMCO29* showed the highest expression at 7d.

These findings indicate that TMCO genes in the resistant HX133 sorghum variety may play crucial roles in response to aphid-induced stress. The differential expression of these genes across time points reflects their specific regulatory functions in aphid stress response mechanisms. This study provides insights into the temporal dynamics of gene expression, supporting the hypothesis that certain TMCO genes contribute significantly to sorghum’s resistance against aphid stress. Further functional validation will help elucidate their precise roles in stress response pathways.

## 4. Discussion

Multicopper oxidases (MCOs) are catalysts of great interest in various biotechnological applications(<u>Valles, Kamaruddin, Wong, & Blanford, 2020</u>), particularly in bioremediation and energy transduction. TMCO proteins play critical roles in multiple aspects of plant biology, including growth and development, responses to biotic and abiotic stresses, and hormone signal transduction. While the biological functions and bioinformatics analyses of laccases and ascorbate oxidases, subfamilies of TMCO, have been extensively studied, much remains unknown about the three-domain multicopper oxidases. Investigating the functions of TMCO gene families in plants can provide valuable insights into plant adaptation mechanisms and help elucidate the role of TMCO genes in aphid resistance in sorghum (*Sorghum bicolor*).

TMCO genes have been identified in many plant species, but detailed analyses have been conducted only in model crops such as rice (*Oryza sativa L.*) and foxtail millet (*Setaria italica*). Research by Chen Huilong et al. revealed that as plants evolved, the number of TMCO family members gradually increased, with more members found in higher angiosperms(<u>Chen Huilong 2021,19(17):5569-5580.(in Chinese)</u>). This suggests an evolutionary expansion of the family, likely driven by whole-genome duplication events, highlighting the potential adaptive significance of these genes.

Most identified TMCO proteins in sorghum are hydrophilic, containing abundant hydrophilic amino acid residues that facilitate hydrogen bonding with water molecules. This characteristic enhances their solubility and stability in aqueous environments, enabling their functional activity in the cytoplasm or extracellular spaces(<u>Damodaran, 2008</u>). Subcellular localization predictions indicate that many TMCO proteins are localized to chloroplasts, vacuoles, and the extracellular matrix, with fewer in the cytosol. The prominent presence in chloroplasts suggests their involvement in efficiently scavenging reactive oxygen species (ROS) under high light conditions, thereby coordinating antioxidative mechanisms across organelles, delaying senescence, and enhancing plant vitality. The instability index of most *SbTMCO* proteins is below 40, indicating relative stability. This property is advantageous for enzyme activity assays and protein-protein interaction studies, further enabling functional exploration of these proteins in stress responses.

Gene duplication plays a critical role in biological evolution, providing a genetic foundation for the emergence of new traits. It is one of the primary driving forces behind the evolution of gene families(<u>Panchy, Lehti-Shiu, & Shiu, 2016</u>). In sorghum (*Sorghum bicolor*), 12 duplicated pairs of TMCO genes have been identified on the chromosomes, of which 8 pairs are tandem duplicates, and the remaining 4 are segmental duplicates. These findings suggest that the duplication and subsequent amplification of TMCO genes in sorghum have occurred throughout its evolutionary history.Calculations of the nonsynonymous (Ka) to synonymous (Ks) substitution rate ratios (Ka/Ks) indicate that all duplicated gene pairs have Ka/Ks values less than 1, implying they are under purifying selection. This suggests that the genes are functionally conserved with a low degree of divergence.

Cis-acting elements play crucial roles as molecular switches in transcriptional regulation(<u>Yamaguchi-Shinozaki & Shinozaki, 2005</u>). The analysis revealed that *SbTMCO* genes are enriched with various hormone-responsive regulatory elements, notably jasmonic acid (JA) and abscisic acid (ABA) response elements. These elements are essential for enabling plants to adapt to and resist environmental stresses by modulating gene expression, synthesizing specific proteins, and integrating hormonal signaling(<u>Zhao, Liu, Wang, & Yuan, 2021</u>). Jasmonic acid (JA), an endogenous hormone, regulates plant growth and development while acting as a signaling molecule in defense responses to both biotic and abiotic stresses(<u>Yu et al., 2019</u>). Studies have demonstrated that methyl jasmonate (MeJA)-treated sorghum seedlings can effectively suppress the infestation of green aphids, underscoring the critical role of JA-mediated defense pathways in aphid resistance(<u>Zhu-Salzman, Salzman, & Koiwa, 2004</u>). Abscisic acid (ABA), another key plant hormone, plays a pivotal role in growth regulation and stress adaptation(<u>Jin, Ni, & Ruan, 2009</u>). Research indicates that ABA significantly contributes to insect resistance, as its deficiency increases plant susceptibility to herbivorous insects(<u>Thaler & Bostock, 2004</u>; <u>Yan et al., 2022</u>). For instance, Esmaeily et al. demonstrated that JA and ABA synergistically enhance tomato resistance to greenhouse whiteflies (Trialeurodes vaporariorum) by inducing defense responses and the production of phenolic compounds(<u>Saeideh, Mohammad, & Hamzeh, 2020</u>). These findings underscore the complex regulatory network underlying plant defenses, where hormones like JA and ABA synergistically regulate adaptive responses to biotic stressors.

In resistant sorghum, several TMCO genes (*SbTMCO36*, *SbTMCO4*, *SbTMCO34*, *SbTMCO37*, *SbTMCO35*, *SbTMCO15*, *SbTMCO29*, *SbTMCO42*, and *SbTMCO31*) exhibit high expression during early stages of aphid infestation (6h, 24h, and 48h). This suggests that these genes may play significant roles in sorghum’s defense against aphids, although their regulatory mechanisms require further investigation.

## 5. Conclusion

This study focused on the sorghum (*Sorghum bicolor*) TMCO gene family, employing bioinformatics tools to analyze the genome-wide distribution of TMCO genes and their differential expression under aphid stress. The main findings are summarized as follows:

1. **Identification and Distribution**: A total of 42 TMCO genes were identified, unevenly distributed across 10 chromosomes. Among them, 8 pairs were involved in tandem duplications, and 4 pairs underwent segmental duplications.
2. **Subcellular Localization and Structural Features**: Subcellular localization predictions indicated that *SbTMCO* proteins are predominantly located in chloroplasts, vacuoles, and the extracellular matrix. Most proteins were unstable in vitro, and their secondary structures were primarily composed of random coils.
3. **Collinearity and Phylogenetic Analysis**: Collinearity and phylogenetic analyses revealed a close evolutionary relationship between sorghum and soybean (*Glycine max*) TMCO genes, with notable collinearity and a shared ancestry between the two species.
4. **Gene Structure and Regulatory Elements**: Analysis of gene structures and conserved motifs showed that genes within the same subgroup exhibited similar exon-intron arrangements. Cis-acting element analysis identified jasmonic acid methyl ester (MeJA)-responsive and abscisic acid (ABA)-responsive elements as the most abundant, suggesting a critical role for TMCO genes in aphid resistance.

### Expression Analysis under Aphid Stress

qPCR analysis demonstrated that nine TMCO genes (*SbTMCO36*, *SbTMCO4*, *SbTMCO34*, *SbTMCO37*, *SbTMCO35*, *SbTMCO15*, *SbTMCO29*, *SbTMCO42*, and *SbTMCO31*) were highly expressed during early stages of aphid infestation (6h, 24h, and 48h), indicating their potential roles in sorghum aphid resistance.

Developing aphid-resistant sorghum varieties remains a significant challenge for breeders(<u>Thudi et al., 2024</u>). Concerns over the excessive use of insecticides and its impact on the environment and grain quality have spurred interest in genetic approaches to pest resistance(<u>Swathy et al., 2024</u>). This study fills gaps in the research on the *SbTMCO* gene family, identifying high-expression genes with potential functional roles in aphid resistance. The findings provide a molecular basis for further investigations into insect resistance in sorghum, offering a valuable genetic resource for breeding efforts.

## Author Contributions

Conceptualization, J.D and D.W.; methodology, J.D.; software, M.G., C.Y. and T.W; validation, T.W., M.G., J.L. and Y.S.; formal analysis, T.W., L.S.; investigation, M.G., T.W. and Y.W.; resources, J.D. and D.W.; data curation, M.G., T.W. and CY.; writing—original draft preparation, M.G., J.L., T.W., Y.S., L.S., Y.W., J.D. and D.W.; writing—review and editing, M.G.,C.Y., Y.S., J.D. and D.W.; visualization, M.G., T.W. and J.L.; supervision, J.D. and D.W.; project administration, M.G.,; funding acquisition, J.D. and D.W. All authors have read and agreed to the published version of the manuscript.

## Ethical approval

Not applicable.

## Informed consent

Not applicable.

